# PlantNetX: A web-based transcriptomic resource integrating bulk and single-cell RNA-seq for plant functional genomics

**DOI:** 10.64898/2026.07.30.741827

**Authors:** Mohsin Ali Nasir, Samia Nawaz, Ahmed Faik

**Affiliations:** Environmental and Plant Biology Department, Ohio University, Athens, Ohio, 45701 USA; Interdisciplinary Program in Molecular and Cellular Biology, Ohio University, Athens, Ohio 45701 USA

**Keywords:** Bulk RNA-seq, co-expression analysis, functional genomics, gene association network, PlantNetX, rice (*Oryza sativa*), single-cell transcriptomics

## Abstract

Bulk RNA sequencing and single-cell RNA sequencing provide complementary information on tissue and cell-type-specific gene expression. Bulk RNA sequencing enables the construction of gene association networks that identify co-expressed genes involved in shared pathways, whereas single-cell RNA sequencing maps their expression to cell types. However, most platforms provide access to either bulk RNA sequencing or single-cell RNA sequencing analysis, making it difficult to connect tissue-level co-expression with cell-type-specific expression. PlantNetX was developed as a web-based platform that integrates both data types. Although PlantNetX currently focuses on rice (Oryza sativa) and includes 70 quality-controlled RNA sequencing datasets comprising 1,198 sequencing libraries, together with nine single-cell RNA sequencing datasets containing more than 580,000 cells, including recently released datasets not consistently represented in existing platforms, it was designed to incorporate additional plant species, datasets, and analytical tools. PlantNetX provides Mutual Rank-based co-expression analysis, global and tissue-specific gene association networks, interactive visualization, and cell-type-specific expression summaries. The platform was validated with published examples of plant cell-wall biosynthesis and root-hair growth and retrieved gene association and expression patterns. Under standardized testing conditions, PlantNetX had a shorter mean response time than the other databases assessed. PlantNetX will support research in plant cell-wall biosynthesis, pathway discovery, functional genomics, and crop improvement.

**Highlights:** PlantNetX integrates gene co-expression networks with cell-type expression, enabling fast identification and biological interpretation of candidate genes across tissues and individual cells.

**Graphical Abstract:** 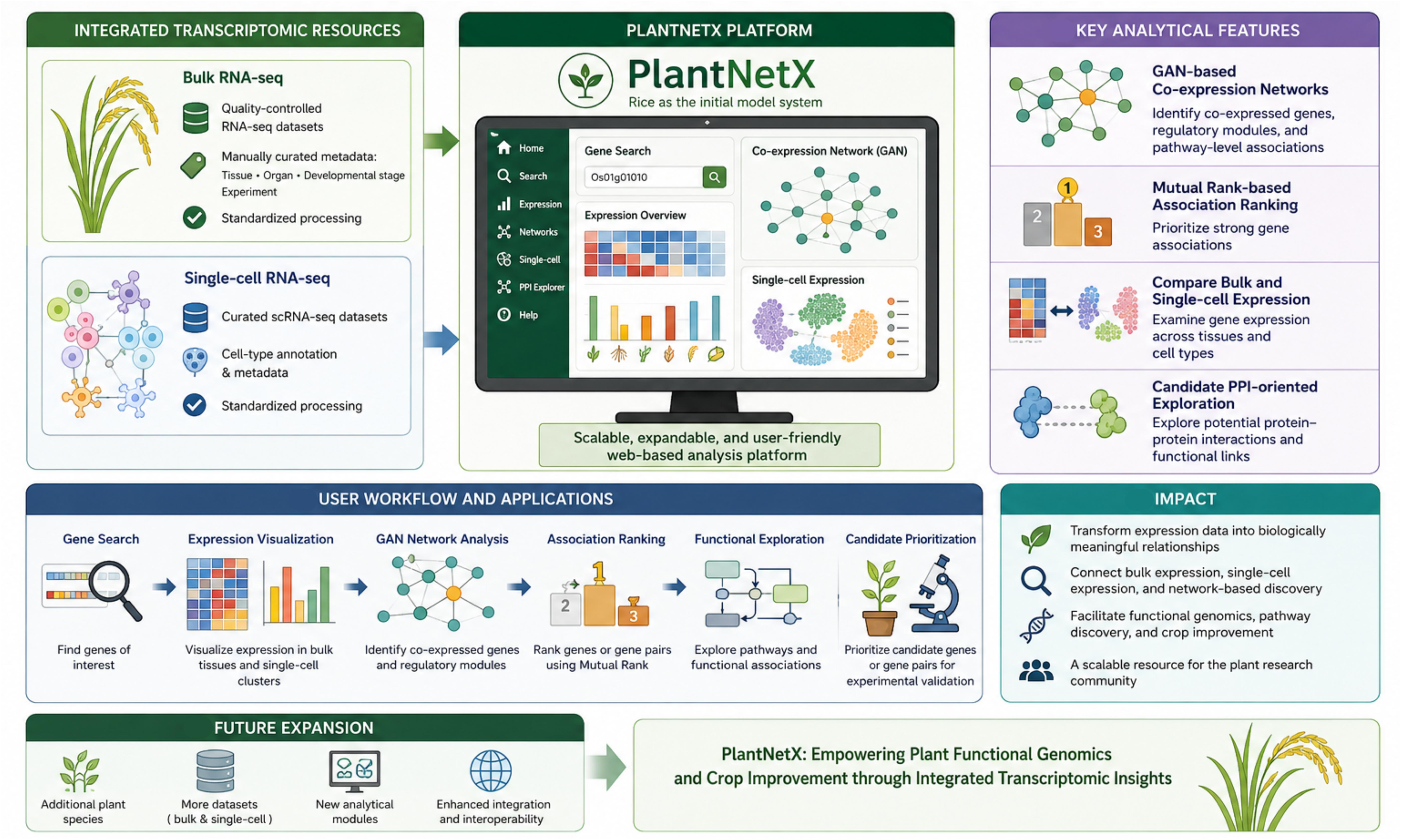

## Introduction

The recent progress in high-throughput sequencing technologies and the development of advanced analytical tools allows an unprecedented understanding of gene expression dynamics and regulatory networks. Specifically, global co-expression profiles from many tissues and organs can be generated from RNA sequencing (RNA-seq) datasets. RNA-seq technologies provide higher sensitivity, larger dynamic range and more accurate quantifications compared to microarray-based technologies (Han *et al*., 2015). The use of these transcriptomic datasets should speed up the field of crop improvement, as many complex agronomic features are controlled by multiple genes that work together in a coordinated manner. Classical methods, such as map-based cloning, quantitative trait locus mapping and genome-wide association studies, have been instrumental in identifying genes associated with important agronomic traits (i.e., yield, stress tolerance, plant architecture, flower development, grain quality, and nutrient use efficiency) (Li *et al*., 2018). However, these methods are labour-intensive and not efficient in correlating the impact of multiple genes on complex agronomic traits. More recently, single-cell RNA sequencing (scRNA-seq) has enabled the addition of a new domain of plant transcriptomics that allows researchers to perform expression analysis on various cell populations within plant organs/tissues. scRNA-seq technology provides high-resolution measurements of gene expression profiles of individual cells in a population, which helps determine cell identity and development trajectories in a plant (Denyer *et al*., 2019; Ryu *et al*., 2019; Zhang *et al*., 2021). While RNA-seq provides the overall level of co-expression of genes for a tissue (and/or condition), the scRNA-seq gives the resolution at the cellular level. However, organization and standardization of scRNA-seq datasets are critical and require development of plant cell atlases including marker genes and gene expression patterns across multiple plant species. For example, the scPlantDB database (He *et al*., 2024) could integrate single-cell transcriptomic profiles from multiple high-quality datasets across 17 plant species. Although these large transcriptomic datasets (RNA-seq and scRNA-seq) can be accessed via public repositories such as the Gene Expression Omnibus (GEO) and Sequence Read Archive (SRA), each dataset requires the performance of quality control, normalization, metadata curation, and computation to be useful to the scientific community (Barrett *et al*., 2012; Han *et al*., 2015; Leinonen *et al*., 2010).

Valuable information can be extracted from Gene-Association Network (GAN) analyses, including identification of gene modules and transcriptional programs that regulate the coordinated expression of multiple genes in a complex biological process (Shang *et al*., 2020; Zhang *et al*., 2022). In these networks, genes are represented by nodes that are connected by edges based on their similarity of expression patterns across a set of biological samples, tissues or experimental circumstances (Ruan *et al*., 2010; Serin *et al*., 2016). Because the similarity in expression patterns tends to reflect contribution to similar biological processes, it is possible to predict the biological function of specific genes and better our understanding of highly complex plant traits (Allocco *et al*., 2004; Van Dam *et al*., 2018). Co-expression analysis has been used to prioritize candidate genes involved in plant cell-wall polysaccharide biosynthesis (Oh *et al*., 2013). To perform accurate and reliable GAN analysis, one needs high-quality transcriptome datasets, consistent processing and normalization, well-curated metadata, and robust methods to measure co-expression correlations. The expression similarity between genes is usually assessed through the Pearson correlation coefficient (PCC) and Mutual Rank (MR). MR prioritizes reciprocal co-expression relationships, helping to identify candidate genes that may participate in shared biological processes or pathways. Lower MR values indicate stronger reciprocal co-expression relationships.

Several plant transcriptome platforms are publicly available for RNA-seq- and microarray-based datasets, including RiceFREND (Sato *et al*., 2013), Rice Expression Database (RED) (Xia *et al*., 2017), OryzaExpress (Hamada *et al*., 2011), PlantExp (Liu *et al*., 2023), and PPRD (Yu *et al*., 2022). But the existing technologies are still limited in some aspects. Some resources are mostly based on older microarray datasets, which have lesser sensitivity and smaller dynamic range compared to RNA-seq. Others provide static or pre-defined co-expression networks, limiting flexible tissue-specific analysis and user-driven network exploration. Moreover, gene searching might be challenging with old gene identities, limited search options and poor association with recent single-cell RNA-seq data sets. Plant single-cell RNA-seq resources have been developed as well, such as PlantscRNAdb (Chen *et al*., 2021), Plant Single Cell Transcriptome Hub (PsctH) (Xu *et al*., 2021), and scPlantDB (He *et al*., 2024). These databases provide important information on plant cell types, marker genes, single-cell expression patterns and visualization of cellular heterogeneity. However, they are mainly meant for single-cell data exploration, marker lookup or cell-type comparison and they don’t allow direct analysis gene expression in relation to GAN analysis from RNA-seq. The major limitation is that the tools to analyze RNA-seq and scRNA-seq datasets are different and housed in different platforms/websites. Thus, researchers are forced to switch between different platforms to determine the relationship between tissue level expression, co-expression interactions and cell type specific expression patterns, which can be challenging considering difference in methods used by these platforms. PlantNetX integrates these capabilities into a single workflow by combining RNA-seq expression, tissue-specific GANs, MR-ranked associations, finding gene partners for subsequent experimental validation and single cell expression. The comprehensive comparison is provided in Table 1.

**Table 1.**
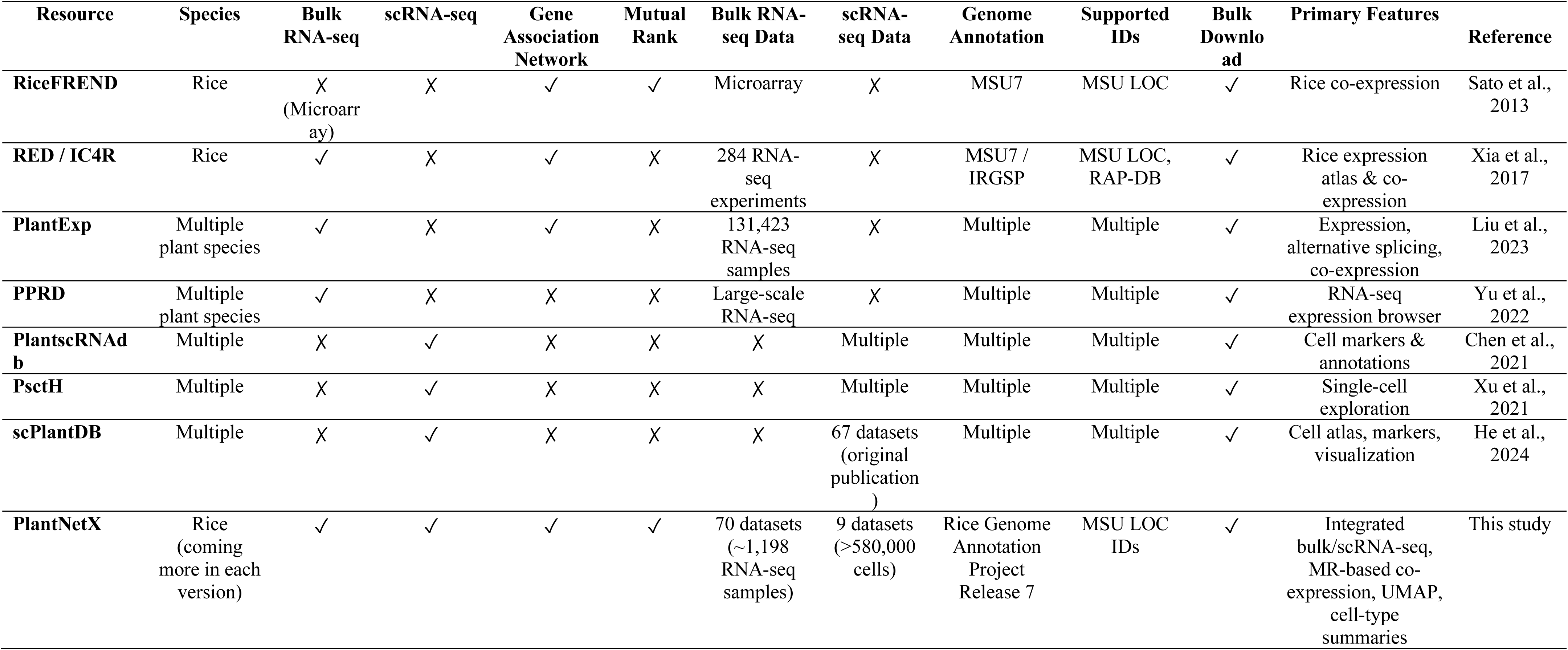
Comparison of PlantNetX with representative plant transcriptomic and single-cell databases. PlantNetX and existing resources were compared using objective criteria, including species coverage, transcriptomic data type, co-expression functionality, MR analysis, dataset scale, genome annotation, supported identifiers, download availability, and primary features. PlantNetX uniquely combines bulk RNA-seq-based co-expression analysis with scRNA-seq-based cell-type expression within one platform.

To overcome these limitations, PlantNetX platform (available at https://plantnetx.academic.kube.ohio.edu/) was developed to offer a single stop for RNA-seq and scRNA-seq analyses. In its current version, the platform is focused on rice (*Oryza sativa*) using 70 quality-controlled RNA-seq datasets (downloaded raw data were manually processed to generate metadata files for tissue types, organ types, developmental stage and experimental conditions) and 9 scRNA-seq datasets. In addition, PlantNetX platform is created with a vision to include other plant models such as Arabidopsis, cotton, and sorghum. In PlantNetX platform, the RNA-seq datasets allows the construction of global and tissue-specific GANs to identify genes with strong co-expression association. These genes can be examined at the level of cell types using scRNA-seq datasets to determine whether certain genes are broadly expressed across a tissue or have a more restricted expression within certain types of cells. Thus, PlantNetX is a valuable genomics resource that is currently needed for progress in crop improvement, pathway discovery and plant functional genomics.

## Materials and Methods

### Data sources

Bulk rice RNA-seq datasets were obtained from the NCBI GEO (https://www.ncbi.nlm.nih.gov/geo) and SRA (https://www.ncbi.nlm.nih.gov/sra) on 10 January 2025. Approximately 100 publicly available rice RNA-seq datasets were initially screened. In this study, a dataset refers to a publication- or project-level accession, whereas a sample refers to an individual sequencing library within that study. Following study- and sample-level quality assessment, 70 datasets containing ∼1,198 sequencing libraries were retained for downstream expression analysis and GAN construction.

These datasets contain transcriptome profiles from different rice tissues such as floret, root, leaf, embryo, grain and seed, which can provide a solid basis for downstream expression analysis and GAN development.

For the scRNA-seq part, PlantNetX has 9 publicly available rice scRNA-seq datasets derived from GEO and SRA. These files include >580,000 cells from various rice tissues and experimental situations. They were selected among 12 accessible rice scRNA-seq datasets regarding data availability, tissue representation, sample information and suitability for downstream single-cell analysis.

Metadata for RNA-seq and scRNA-seq datasets were manually annotated to standardize information on accession numbers, tissue or organ source, developmental stage, experimental conditions, paper description, and dataset provenance. This manual curation was required due to differences in sample names, tissue labels, completeness of metadata and depth of annotation across available datasets.

### Data processing

PlantNetX source code, dataset metadata, database files, and supporting resources are publicly available through the PlantNetX GitHub repository at https://github.com/Mohsin-OU/PlantNetx. The PlantNetX web resource is publicly accessible at https://plantnetx.academic.kube.ohio.edu/.

Raw RNA-seq data were downloaded from the NCBI SRA (Leinonen *et al*., 2010) and converted to FASTQ format using fastq-dump software of SRA Toolkit v2.8.2. All data sets were paired-end sequencing libraries. Read quality and adapter sequences were assessed using FastQC (Andrews, 2010) v0.11.5 and low-quality bases were trimmed using fastp v0.23.4 with custom parameters (Chen *et al*., 2018). Samples of poor overall sequencing quality or with an excess of over-represented sequences were excluded from further analysis.

High quality reads were mapped to the Oryza sativa reference genome acquired from the Rice Genome Annotation Project Release 7 (http://rice.plantbiology.msu.edu/, accessed on 13 February 2025) after trimming. Alignment was done with STAR v2.5.3 using options --quantMode GeneCounts, --outSAMtype None, --outSAMmode None and -- readFilesCommand zcat (Dobin *et al*., 2013). Genome indices were constructed with sjdbOverhang=100, genomeChrBinNbits =14, genomeSAindexNbases=12 and genomeSAsparseD=3. The strandedness of the library was inferred from the metadata of each dataset and the appropriate STAR gene-count output was utilized to quantify the expression.

Gene-level read counts generated by STAR were translated to transcripts per million (TPM) using a customized Python script gene_length.py. TPM calculations were adjusted to include gene lengths from the Rice Genome Annotation Project GTF annotation to account for differences in gene length and library size (Wagner *et al*., 2012). TPM values were processed into log₂ (TPM + 1) for downstream analysis.

Samples were retained for downstream analysis only if they met all of the following criteria: (i) at least 70% of reads remained after adapter and quality trimming, (ii) more than 70% of reads mapped uniquely to the rice reference genome, and (iii) at least 10 million uniquely mapped reads were obtained. Following quality verification, a total of 70 high-quality rice RNA-seq studies were selected for the development of the GAN.

TPM normalization accounts for gene length and library size within each sample, which increases the comparability of gene expression estimations among RNA-seq datasets obtained independently. The log_2_ (TPM+1) transformation was used to stabilize variance, limit the effect of highly expressed genes and enhance the robustness of the co-expression analysis based on the PCC.

Since PlantNetX uses RNA-seq datasets collected by several studies and different experimental conditions, some residual biological and technical variation is expected. This batch-to-batch variance is typical in large-scale transcriptome datasets and can influence downstream studies (Leek *et al*., 2010). To minimize these effects, all datasets were analyzed with the same analysis pipeline including uniform quality assessment, read trimming, genome alignment, gene quantification and TPM normalization following standard RNA-seq analysis techniques (Conesa *et al*., 2016). Prior to network development, genes with TPM <1 were excluded to reduce the contribution of low-abundance transcripts, which may introduce sampling noise into downstream expression analyses (Sha *et al*., 2015) and affect the reliability of downstream expression assessments. Pearson correlation analysis was conducted with repeated resampling to evaluate the consistency of gene–gene associations and increase the stability of the co-expression estimations. Resampling and bootstrap-based methods are often used to assess resilience and repeatability of inferred biological networks. The raw read counts at the gene level obtained by STAR were converted to TPM using the conventional normalization approach to account for gene length and library size (Wagner *et al*., 2012) as follows:

Reads Per Kilobase (RPK) was calculated as Equation 1:

where:

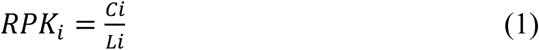

- *C_i_*= raw read count of gene *i* obtained from STAR GeneCounts
- *L_i_*= gene length of gene *i* (in kilobases) TPM was calculated using Equation 2.

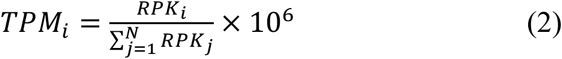

where *C_i_* denotes the raw read count of gene *i*obtained from STAR GeneCounts, *L_i_* denotes the gene length (in kilobases) extracted from the Rice Genome Annotation Project Gene Transfer Format (GTF), *RPK_i_* represents the RPK value of gene *i*, and *N* denotes the total number of annotated genes in the sample. TPM normalization accounts for both gene length and sequencing depth, thereby facilitating more reliable comparisons of gene expression across independently generated RNA-seq datasets.

For scRNA-seq analysis, raw FASTQ data were quality checked and adaptor sequences and low quality bases were trimmed using fastp. High-quality reads were mapped to the rice reference genome, adjusted for cell barcodes, processed for unique molecular identifiers (UMIs) and gene-by-cell count matrices were created using STARsolo (Kaminow *et al*., 2021). Downstream analyses were performed using a Scanpy-based workflow (Wolf *et al*., 2018) that included quality filtering, library-size normalization, logarithmic transformation, identification of highly variable genes, principal component analysis (PCA), Leiden clustering (Traag *et al*., 2019), and Uniform Manifold Approximation and Projection (UMAP) for dimensionality reduction (McInnes *et al*., 2018). Doublets were predicted and removed using DoubletFinder (McGinnis *et al*., 2019) prior to further analyses. To reduce technical variation across independently generated scRNA-seq datasets, batch correction and dataset integration was performed using the Seurat integration method (Stuart *et al*., 2019). This approach permits alignment of shared biological cell types across studies, while minimizing technical variation arising from differences in library preparation, sequencing platform, sequencing depth, and study-specific processing.

Cell-type annotations were first obtained from the original studies and then checked and harmonized based on published rice cell-type marker genes and PlantscRNAdb. Marker gene evidence was used to validate and refine cell-type assignments where necessary, thereby improving annotation consistency across datasets. After quality control, doublet removal, and Seurat-based integration, the scRNA-seq datasets were combined into a unified analysis framework and visualized in a global UMAP embedding. The shared UMAP coordinates were used to represent the integrated cellular landscape across datasets and to compare gene expression among annotated cell types. This strategy maintains biological structure within individual datasets while limiting the effect of technological variance on downstream visualization and interpretation. The processed data matrices were recorded as AnnData objects (.h5ad) with normalized expression matrices, cell metadata, cluster assignment, UMAP coordinates, and annotated cell types. This standardized framework allows to do efficient gene-level queries, cell-type-specific expression summary and interactive visualization in PlantNetX.

### GAN Construction

A GAN and tissue-specific subnetworks were built using normalized RNA-seq expression datasets in PlantNetX for co-expression calculation. To minimize noise from very lowly expressed genes, only the genes with TPM ≥ 1 were kept in the dataset group being investigated for co-expression computation. For the respective analysis, genes with TPM < 1 were excluded from the PCC and MR-based network design, as very low expression values may result in unstable or biologically weak relationships. PlantNetX also curates expression data for all genes in the database, so users can obtain gene-level expression information even when a gene is not part of a given network calculation. Thus, users can view the expression profiles of these genes.

Correlation of the expression of genes was evaluated using PCC for each gene pair using the following Equation 3:

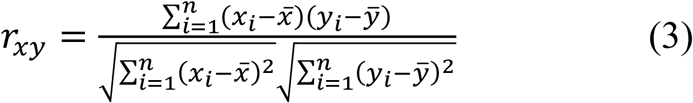

where x_1_ and y_1_ are the log₂ (TPM + 1) expression values of the two genes in sample i, x̄ and ȳ are their mean expression values across all samples, and n is the total number of samples included in the analysis. Pearson correlation was calculated without sample-specific weights.

To assess the consistency of gene–gene correlations, PCC values were calculated across 1,000 resampling iterations. In each iteration, 80% of the retained RNA-seq samples were selected at random without replacement, and PCC values were recalculated for all gene pairs. The final PCC for each pair was obtained by averaging the correlation coefficients across iterations. Fisher z-transformation was not applied before averaging. Edge stability was not evaluated as a separate metric, and no explicit tissue-balancing procedure was applied during construction of the global network. To provide a more biologically focused view and reduce the influence of uneven tissue representation, PlantNetX also includes tissue-specific subnetworks that allow co-expression relationships to be examined within individual tissue groups.

Gene pairs were ranked according to their MR values, and lower MR values were considered to represent stronger and more reliable reciprocal co-expression relationships.

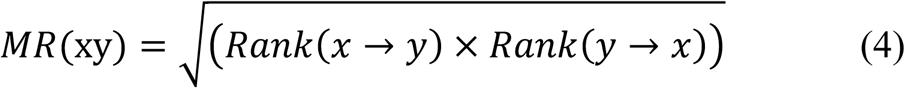

Here in Equation 4, *Rank*(*x* → *y*)denotes the rank of gene *y*among all genes co-expressed with gene *x*, whereas *Rank* (*y* → *x*)denotes the reciprocal rank of gene *x* among all genes co-expressed with gene *y* . Lower MR values indicate stronger reciprocal co-expression relationships and therefore provide greater confidence in the inferred co-expression between two genes.

### Cell-type Annotation and Marker Information

Cell type annotation of the PlantNetX single cell datasets was performed on the basis of original dataset annotation, cluster-specific expression patterns, tissue origin and marker-supported interpretation. Since the scRNA-seq datasets used in PlantNetX were obtained from publicly available studies, the original cell-type labels were kept when available. These labels were manually reviewed and coordinated across datasets to reduce discrepancies in naming style. Where original labels were missing, partial, or conflicting, broader cell-type identities were inferred from cluster-level expression patterns and tissue of origin. The marker information was obtained from original dataset annotations, published rice scRNA-seq research, and PlantscRNAdb (http://ibi.zju.edu.cn/plantscrnadb/%/celltype), a publicly available plant single-cell marker and cell-type database (Chen *et al*., 2021). Cluster-specific marker genes together with their differential expression statistics, enrichment scores, and corresponding annotated cell types are provided in in the PlantNetX GitHub repository at https://github.com/Mohsin-OU/PlantNetx.

While reference and marker-based approaches are effective for assigning known cell identities, they can be influenced by variations in tissue source, sequencing platform, preprocessing pipeline, clustering resolution and reference quality. Ambiguous, transitional, or unusual cell populations may be forced into the closest known label if the cell state is not in the reference. The PlantNetX approach considers marker information as complementary evidence rather than a proof. Therefore, cell type labels were finalized based on a combination of marker support, original annotation, tissue origin and cluster-level expression patterns. Cluster-level annotation was informed by marker enrichment. For a cluster “c” and candidate cell type “t”, the marker score was computed as Equation 5:

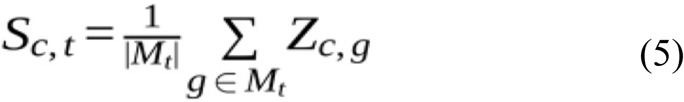

where S (c, t) is the annotation score of cluster “c” and candidate cell type “t”, M_t is the marker gene set of cell type “t”, and Z_(c,g) is the scaled average expression of marker gene “g” in cluster “c”. The scaled expression value is computed as Equation 6:

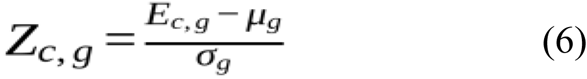

where E (c, g) is the mean normalized expression of gene “g” in cluster “c”, mu_g is the mean expression of gene “g” for all clusters, and sigma_g is the standard deviation of gene “g” for all clusters. The marker score was used as a quantitative measure supporting cell-type assignment. Final annotations were assigned to the candidate cell type exhibiting the highest marker score together with consideration of tissue origin, original study annotations, and biological context.

PlantNetX summarizes gene expression for each annotated cell type by reporting mean expression, the percentage of cells with detectable expression, the number of expressing cells, and the total cell count. The percentage of cells that express a gene in each cell type was computed as Equation 7:

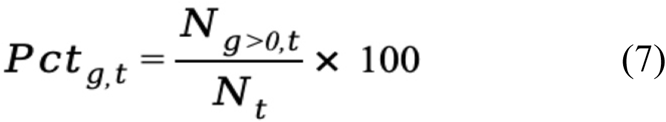

where N (g > 0, t) is the number of cells in cell type “t” with detectable expression of gene “g” and Nt is the total number of cells assigned to that cell type.

The average gene expression within a cell type was determined as Equation 8:

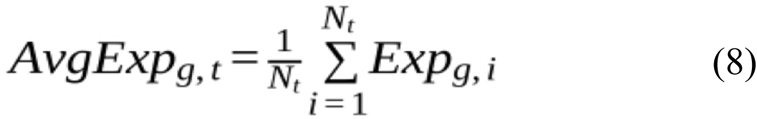

where Exp (g, i) is the normalized value of expression of gene “g” in cell “I”, N t is the total number of cells in the cell type. These values help users to know whether a gene has broad expression across cell types or enrichment in a particular cell population.

### Data Implementation and Database Development

The PlantNetX is hosted in a server having a backend that was built with the Django 3.1 framework (https://www.djangoproject.com/), Python 3.11., and PostgreSQL (https://www.postgresql.org/) was used to create the database. The frontend of the server was created using HyperText Markup Language (HTML), Cascading Style Sheets (CSS) and JavaScript. All requests from the frontend to the database are made asynchronously using asynchronous JavaScript and XML (AJAX) and the data is sent in JavaScript Object Notation (JSON) format. This method removes the need to reload the page each time, and allows to change the content dynamically, making it more interactive and responsive. The database holds RNA-seq data with gene annotation, normalized expression, curated metadata, tissue categories, PCC scores, MR values and ranked co-expression partners.

For the single-cell component, .h5ad files were processed and linked to the backend database. Dataset metadata, cell annotations, cluster labels, UMAP coordinates, and gene expression summaries were systematically integrated into a unified framework to enable gene-level querying and cell-type-specific downstream analyses. The backend fetches only the appropriate gene expression information or cell-level subset per query, avoiding loading entire single-cell datasets into memory unnecessarily. This solution is based on the single-cell design of PlantNetX, in which the backend organizes the metadata of dataset, cell-level annotation, matrix of gene expression, and pre-computed UMAP embedding for rapid query, dataset filtering, and quick gene lookup.

For the display of networks, PlantNetX includes Cytoscape.js, a general-purpose JavaScript toolkit for the visualization and analysis of biological networks (Franz *et al*., 2016). The solution enables zooming, panning, node-edge highlighting, and several layout variations to satisfy various needs of network research.

PlantNetX calculates the expression of the query gene on the precomputed two dimensional UMAP coordinates for single cell display. Expression intensity is the normalized, log-transformed UMI expression. Cells with an expression value greater than 0.1 were categorized as expressing cells, and cells with values less than 0.1 were considered as non-expressing. The color scale indicates the relative expression level of the query gene in individual cells and enables users to detect cluster-specific or cell-type-enriched expression patterns. Cell-type summary tables show the average normalized UMI expression, percentage of cells expressing, number of cells assigned to the cell type and total number of cells analyzed.

The PlantNetX platform and all its analytical outputs are freely available without registration requirements at https://plantnetx.academic.kube.ohio.edu/. PlantNetX interface supports single-gene, multiple-gene, gene annotation, RNA-seq expression visualization, GAN-based network output, MR-based co-expression ranking, single-cell UMAP visualization, cell-type expression summary and downloadable result tables. This allows users to go from a gene search to tissue level expression, network-based association and cell-type level interpretation all in the same platform.

PlantNetX is hosted on the computational infrastructure of the Ohio Supercomputer Center and maintained in collaboration with Ohio University OIT. The database is reviewed periodically for the addition of newly published rice bulk RNA-seq and scRNA-seq datasets, updates to genome annotation, and improvements in gene functional annotation. Each public release will be accompanied by a version number, release date, and updated metadata to support reproducibility. The web interface has been tested using current versions of Google Chrome, Mozilla Firefox, Microsoft Edge, and Apple Safari. Computationally intensive analyses are performed offline during database construction, while the web platform provides access to precomputed expression profiles, PCC and MR results, co-expression networks, and single-cell summaries. Interactive network performance depends on the number of nodes and edges displayed, as well as the user’s browser and available system memory. Large networks and processed datasets can therefore be downloaded for external analysis. Processed expression matrices, curated metadata, PCC and MR tables, marker-gene information, and single-cell expression summaries are available through the PlantNetX Download page and the associated GitHub repository at https://github.com/Mohsin-OU/PlantNetx.

### Response-time benchmarking

The response time of PlantNetX was compared with that of representative plant transcriptomic and single-cell databases, including RiceFREND, RED/IC4R, PlantExp, PPRD, PlantscRNAdb, PsctH, and scPlantDB. For each platform, a standardized gene query was submitted, and the time from query submission to full display of the principal results page was recorded. Each database was tested in 20 independent runs using the same computer, browser, internet connection, query gene, and testing period. The mean response time was calculated across replicate measurements. Only successful queries that returned the expected output were included in the analysis.

## Results

### Architecture of PlantNetX platform

PlantNetX was designed as a centralized web-based platform to explore plant transcriptome data through the RNA-seq and the scRNA-seq datasets using rice as the first model system. The website is arranged in a friendly-user way that guides users from basic database access to gene level analysis, tissue level expression, gene relationship networks and cell type level expression interpretation. The homepage offers access to two major analytical sections: RNA-seq Data Analysis and Single Cell Data Analysis shown in Fig. 1A. The RNA-seq section consists of bulk transcriptomic expression, functional annotation and GAN analysis (Fig. 1B). The single-cell section contains visualization using UMAP (Fig. 1C), expression summary by cell type, and investigation of gene expression at cellular resolution.

**Fig. 1.**
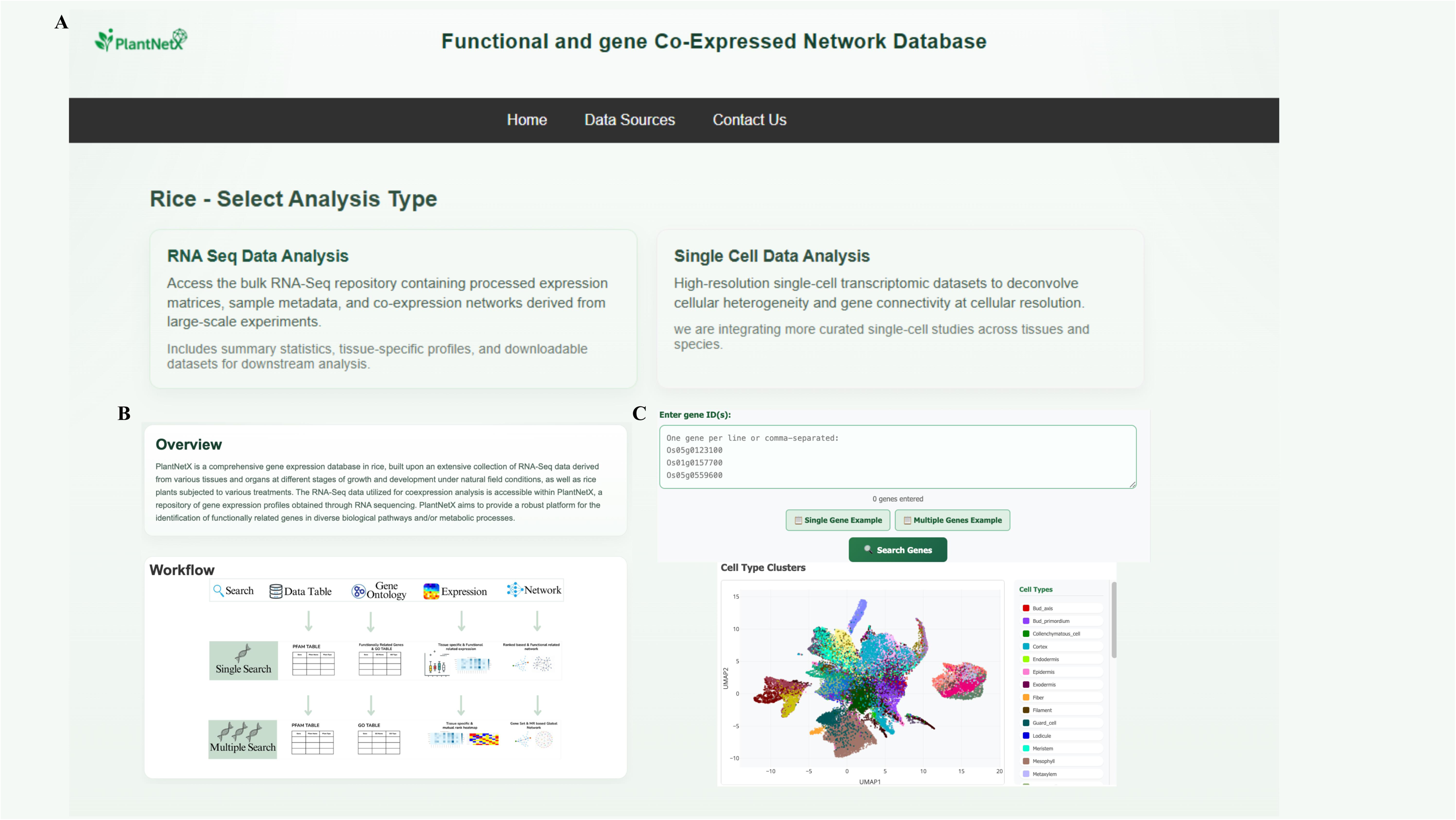
PlantNetX overview and architecture. (A) PlantNetX home page with access points to RNA-seq Data Analysis and Single Cell Data Analysis. (B) RNA-seq module showing gene search, expression display, GO and network functionality. (C) Single cell module including gene search, UMAP display, cell type filtering, summary statistics, and cell type expression information. Supplementary materials include the user handbook and comprehensive workflow views. **ALT TEXT:** Three-panel overview of the PlantNetX platform. The home page provides access to bulk RNA-seq and single-cell analysis modules. The bulk RNA-seq section includes gene search, expression, annotation, and network analysis, while the single-cell section includes gene search, UMAP visualization, cell-type filters, and expression summaries.

#### RNA-seq datasets

The PlantNetX RNA-seq section contains 70 quality-controlled rice RNA-seq datasets comprising approximately 1,198 sequencing libraries (Supplementary Table S1) after quality assessment and filtering to capture global gene expression and GAN patterns. Metadata were manually annotated into major tissue types to facilitate both global and tissue-specific GAN development. The global GAN provides a bird’s eye perspective of gene correlations in all the curated datasets, whilst tissue-specific GANs allow users to explore co-expression patterns in specific biological contexts. The RNA-seq architecture consists of public data collection, standardized processing, metadata curation, PostgreSQL database storage, backend query handling, and interactive online display (Fig. 2A). The backend includes gene search, expression retrieval, PCC/MR outputs, co-expression queries and network generation. The frontend includes dynamic result tables, tissue-level expression graphs, interactive networks and downloadable outputs. The distribution of the curated datasets demonstrate that the major tissue groupings are seed, root and leaf while other categories such as grain, embryo, anther and others provide additional expression contexts (Fig. 2B). The interface allows you to search for single genes or for several genes. The main outputs are Data Table, Gene Ontology (GO), Expression and Network views. The data Table shows gene identifiers, Pfam information, domain type, E-value and annotations. The Expression view presents boxplots and heatmaps, and the Network view shows MR-ranked co-expression relationships utilizing Cytoscape.js for browser-based visualization (Franz *et al*., 2016).

**Fig. 2.**
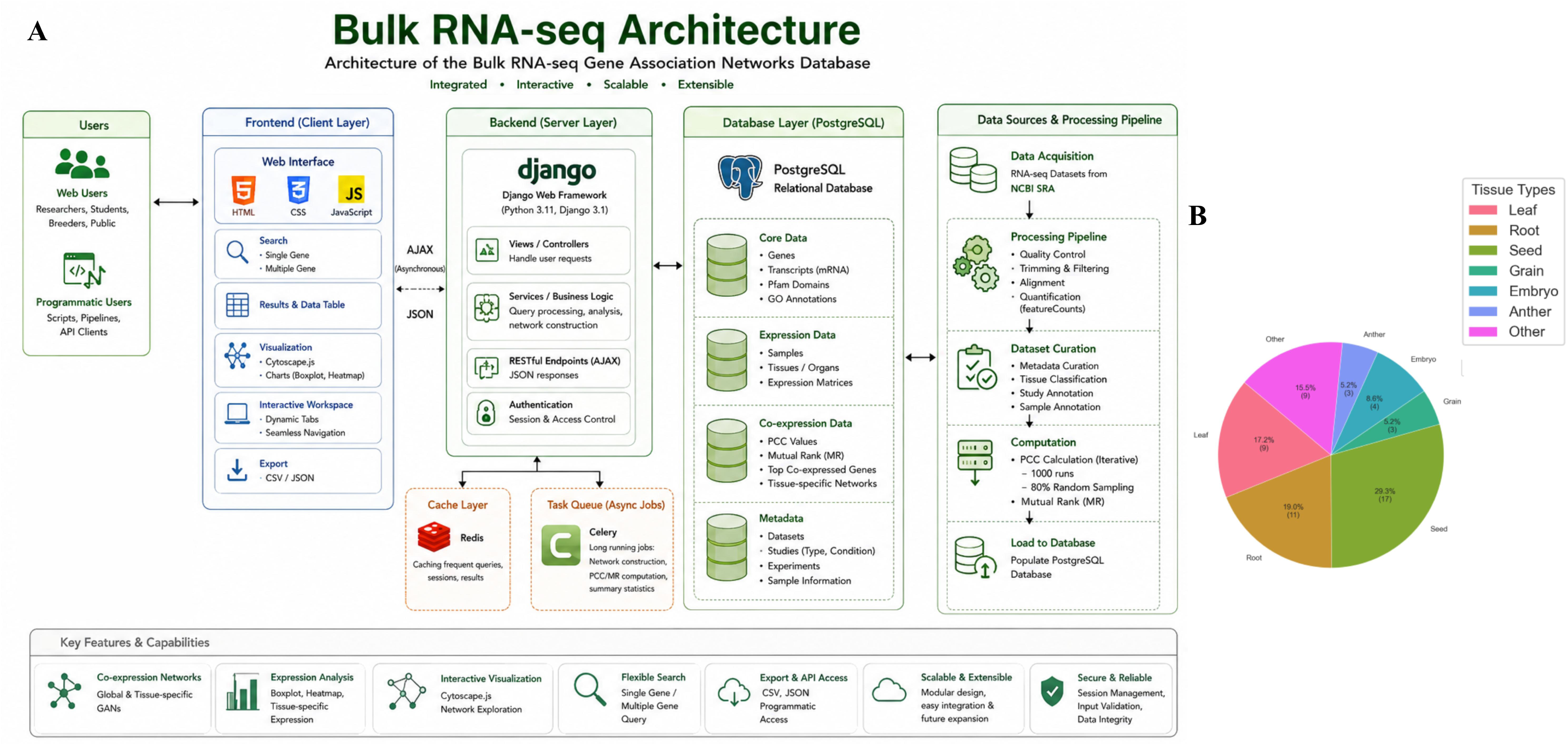
PlantNetX architecture and tissue distribution of bulk RNA-seq. (A) Architecture of the bulk RNA-seq module includes users, frontend layer, backend layer, PostgreSQL database layer, and data sources/processing pipeline. (B) Distribution of curated RNA-seq datasets among main tissue categories. **ALT TEXT:** Two-panel figure showing the architecture of the PlantNetX bulk RNA-seq module and the distribution of curated datasets across rice tissues. The workflow links public RNA-seq data processing, database storage, backend analysis, and web-based visualization. Seed, root, and leaf are the most represented tissue groups.

### scRNA-seq datasets

The PlantNetX single-cell section allows for gene expression analysis at the cellular level. In contrast to the bulk RNA-seq component, which captures gene expression and co-expression relationships at the tissue level, the single-cell component provides cellular-resolution expression profiles through dataset summaries, UMAP visualization, and cell-type-specific expression analysis (Fig. 3). The current single-cell overview consists of 9 datasets (Supplementary Table S2), 81 samples, 586,103 cells, 13 tissues and 39 annotated cell types (Fig. 4A). The tissues are bud, culm, flag leaf and inflorescence, floret, inflorescence, leaf, meristem, pistil, root, root tip, seedlings, shoot and spikelet (Fig. 4B). The section offers a database summary view with count of datasets, count of cells, count of samples, tissue coverage and cell-type coverage. It also offers tissue distribution and dataset contribution charts to display the distribution of the single-cell data in studies and biological sources (Fig.s 4C and 4D). Plots of dataset contribution indicate variation in the number of cells and samples across accessions. For instance, PRJCA004855 provides 303,090 cells and 22 samples, PRJNA974167 provides 116,564 cells and 31 samples. Additional datasets such as PRJNA609100, PRJNA706435, PRJNA663193, PRJNA767589, PRJNA859914, PRJNA922574, and PRJNA1188718 add smaller, but biologically relevant, cell populations. This gives the user an idea of the scale and coverage of each dataset before evaluating gene expression results.

**Fig. 3.**
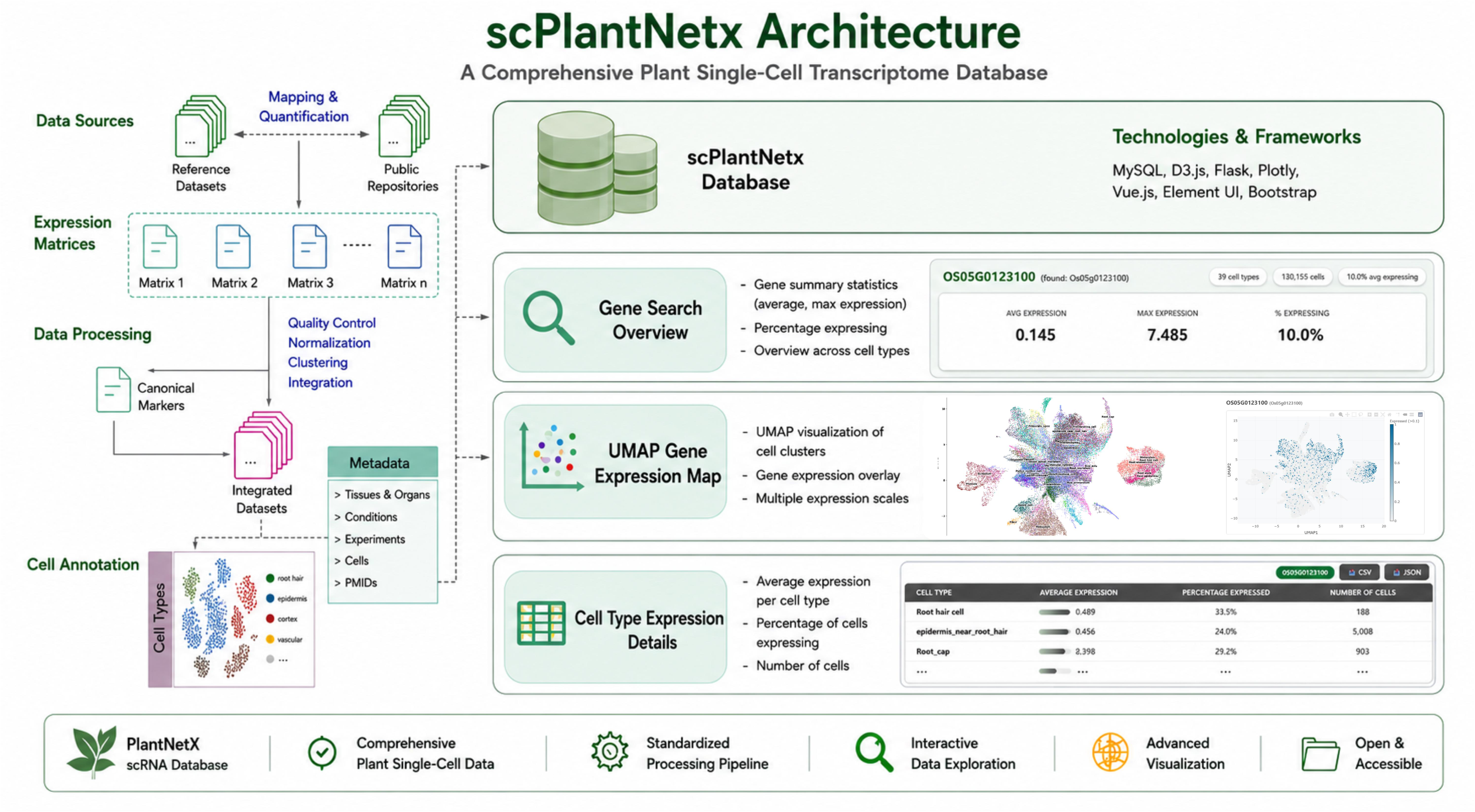
Architecture and interface of the PlantNetX single-cell module. Single cell module architecture: data sources, expression matrices, processing, metadata organization, cell annotation, database integration, gene search overview, UMAP gene expression map, and details of cell-type expression. **ALT TEXT:** Workflow of the PlantNetX single-cell module, showing public dataset collection, processing, cell annotation, database integration, gene search, UMAP visualization, and cell-type-specific expression summaries.

**Fig. 4.**
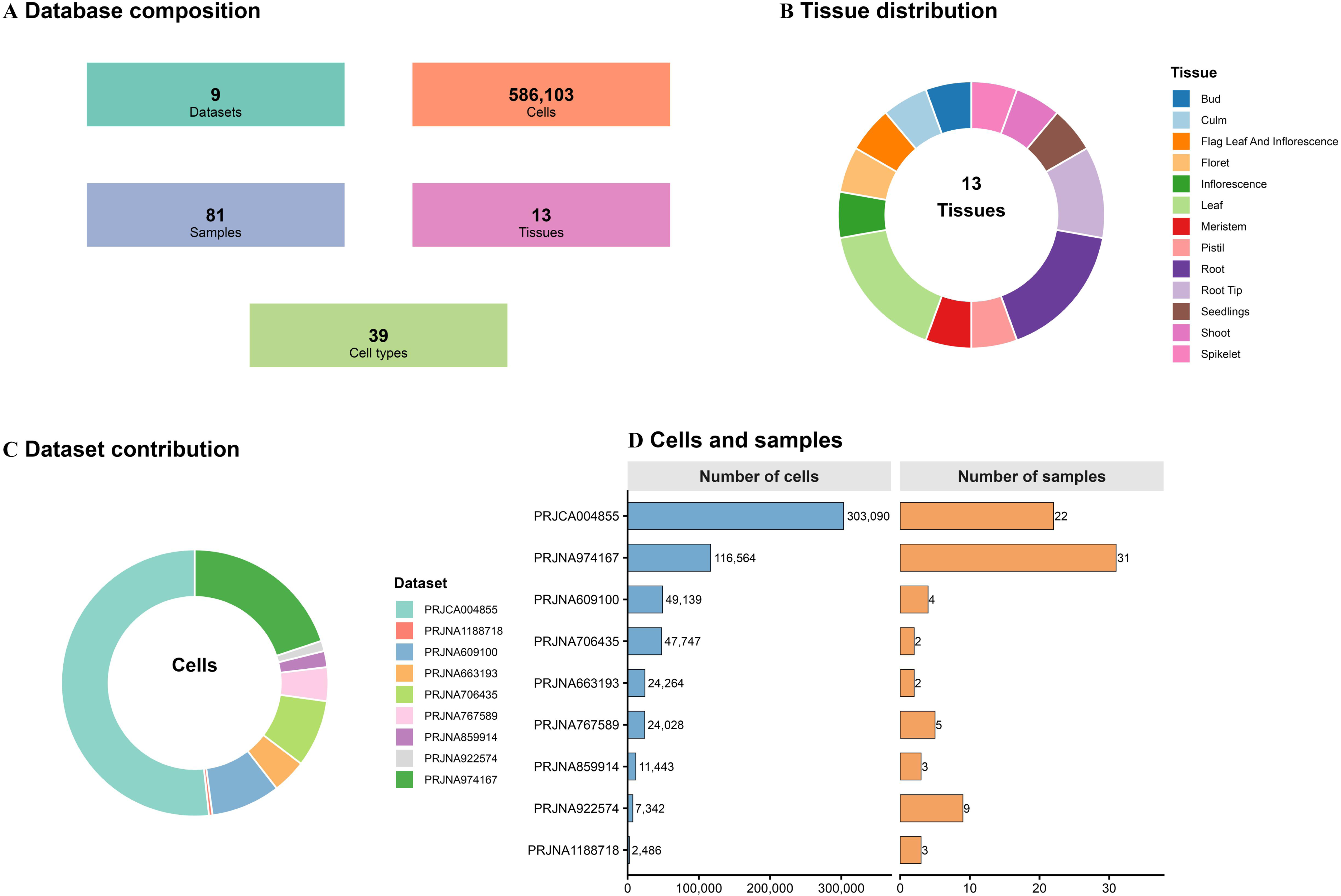
PlantNetX plant single-cell database overview. (A) Composition of the database including number of datasets, cells, samples, tissues and cell types. (B) Tissue distribution of the single-cell collection. (C) Contribution of dataset by cell number. (D) Number of cells and samples provided by each dataset accession. **ALT TEXT:** Four-panel summary of the PlantNetX single-cell database. The database includes nine datasets, 81 samples, 586,103 cells, 13 tissues, and 39 annotated cell types. Additional plots show tissue distribution and the number of cells and samples contributed by each dataset.

### Download and downstream use

PlantNetX outcomes are available for download in comma-separated values (CSV) or JSON formats, including gene annotations, RNA-seq expression profiles, MR-ranked relationships and single-cell expression summaries. These files are useful for pathway enrichment, gene family comparison, candidate gene prioritization, network analysis, figure preparation, and experimental design.

### User Interface for Search and Analysis of RNA-seq datasets

The PlantNetX RNA-seq search interface enables users to explore gene annotation, expression profiles and GAN-based co-expression results in single-gene or multi-gene search modes. PlantNetX currently uses Michigan State University locus identifiers (MSU LOC) identifiers for gene searches. Rice Annotation Project Database (RAP-DB) identifiers can be converted to MSU LOC identifiers using the RAP-DB ID converter https://rapdb.dna.naro.go.jp/converter/. Built-in identifier conversion is planned for a future PlantNetX release.

For single-gene search, users can input one rice gene ID into the search box. An illustration is shown in Fig. 5A, where *LOC_Os01g70190 that is* Glycosyltransferase (GT) 47 was utilized as an example. After submission, PlantNetX sends a gene-specific data table including the gene ID, mRNA ID, Pfam accession, Pfam word, GO information, domain information, protein family annotation and external database linkages (Fig. 5B). The same query also provides tissue-level expression outputs such as boxplots and heatmaps to compare expression patterns across all samples and tissue groupings (Fig.s 5C and 5D). Then users can use/click the co-expression option to see the top genes with MR values for *LOC_Os01g70190* (Fig. 5E). The data are displayed in a table and can be exported as a CSV file for downstream analysis.

**Fig. 5.**
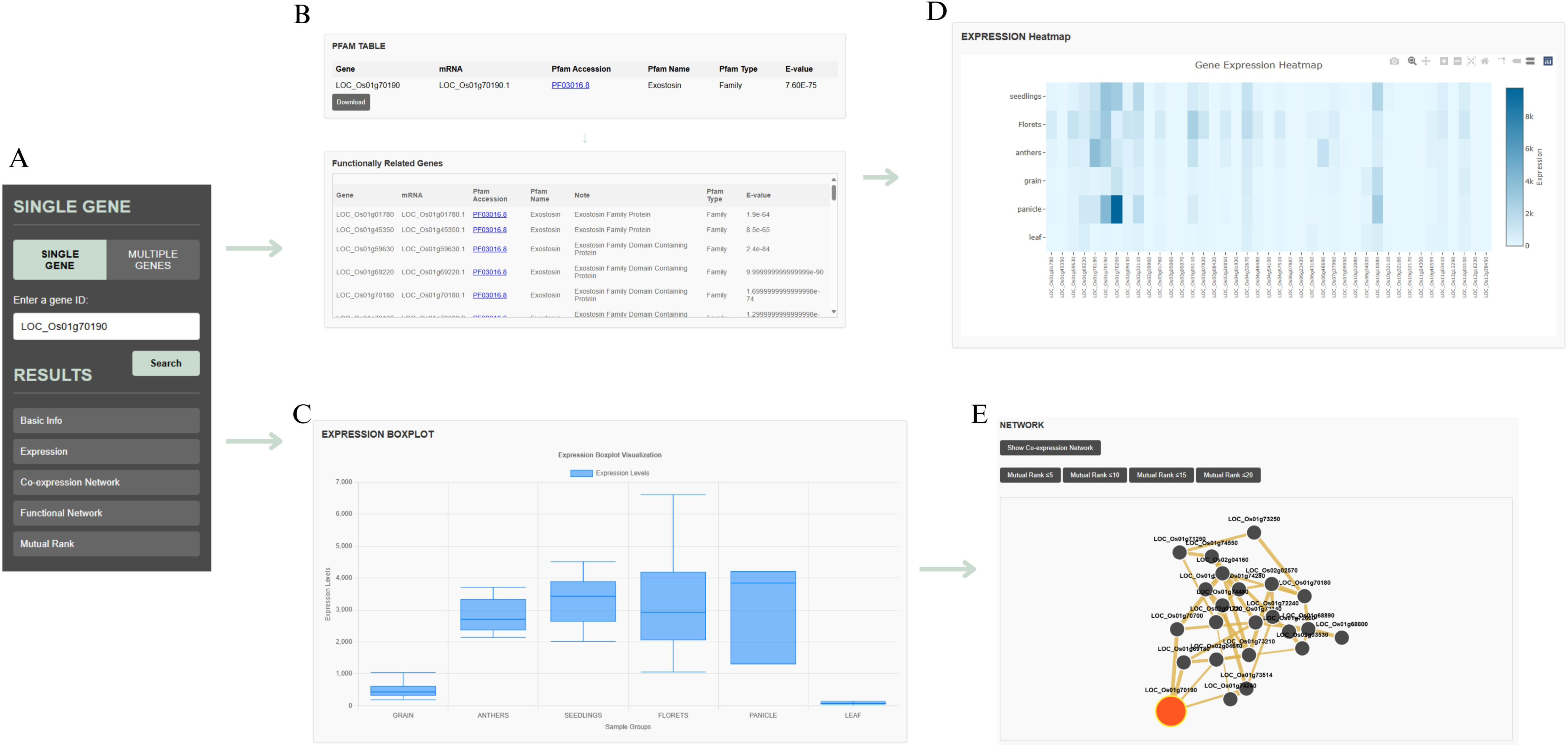
Bulk RNA-seq single-gene search and co-expression analysis in PlantNetX. (A) PlantNetX single-gene query interface showing the input field and available result categories. (B) Gene information for *LOC_Os01g70190,* including the mRNA identifier, Pfam accession, Pfam term, domain type, E-value, and functionally related gene table. (C) Tissue-level expression boxplot for *LOC_Os01g70190* across the major rice tissue groups. (D) Heatmap showing the expression pattern of *LOC_Os01g70190* across the curated RNA-seq samples and tissue categories. Color intensity represents log₂(TPM + 1) expression. (E) Co-expression network showing the top MR-ranked genes associated with *LOC_Os01g70190*. Lower MR values indicate stronger reciprocal co-expression relationships. **ALT TEXT:** Five-panel example of a single-gene search for LOC_Os01g70190 in PlantNetX. The panels show the search interface, gene annotation, tissue-level expression boxplot, expression heatmap, and a co-expression network of the strongest MR-associated genes.

For multiple-gene analysis, the user can input a list of MSU gene IDs in the multiple-gene search box. This tool is helpful for gene family comparison, pathway-level analysis, and validation of experimentally supported candidate gene groups. For illustration, 24 GT genes were used as query/guide genes (Fig. 6A), namely *LOC_Os01g70180, LOC_Os01g70200, LOC_Os01g70190, LOC_Os10g10080, LOC_Os04g32670, LOC_Os03g01760, LOC_Os03g07820, LOC_Os02g32110, LOC_Os05g03174, LOC_Os03g17850, LOC_Os07g49370, LOC_Os01g06450, LOC_Os05g48600, LOC_Os01g48440, LOC_Os10g13810, LOC_Os04g01280, LOC_Os04g55670, LOC_Os06g47340*, *LOC_Os01g54620, LOC_Os10g32980, LOC_Os07g10770, LOC_Os03g62090, LOC_Os07g14850,* and *LOC_Os09g25490*. These genes were selected because it has been shown that they encode for GTs that form specific protein-protein interactions (PPIs) to form several xylan synthase complexes (XSCs) and cellulose synthase complexes in rice (Javaid *et al*., 2024). These genes therefore provide a suitable test set for evaluating whether PlantNetX recovers previously reported co-expression relationships and for identifying candidate gene partners for subsequent experimental validation. When these genes were submitted as query, outcome results were a gene-set annotation table that includes mRNA identifiers, Pfam domain classifications, accession numbers, E-values, and functional information (Fig. 6B). The platform also produces integrated visualization outputs, including a gene network diagram among queried genes (Fig. 6C), tissue-level expression heatmap (Fig. 6D), expanded GAN with additional co-expressed partners (Fig. 6E), and MR heatmap showing pairwise association strength among the provided genes (Fig. 6F). Importantly, PlantNetX recreated similar association patterns among those genes (Fig. 7) similar to published GAN analysis (Javaid et al., 2024). In particular, the reconstructed network consisted of three subnetworks. Subnetwork 1 includes *OsGT47-3 (LOC_Os01g70190), OsGT47-1 (LOC_Os01g70180), OsGT43I (LOC_Os04g55670), OsCesA5 (LOC_Os03g62090)* and *OsCesA6 (LOC_Os07g14850)*. *OsGT47-4 (LOC_Os10g10080), OsGT43J (LOC_Os06g47340), OsGT43B (LOC_Os03g17850), OsGT43F (LOC_Os01g48440), OsGT47-2 (LOC_Os01g70200), OsCesA4 (LOC_Os01g54620), OsCesA7 (LOC_Os10g32980), OsCesA8 (LOC_Os07g10770)* and *OsCesA9 (LOC_Os09g25490*) were in subnetwork 2. *OsGT43G (LOC_Os10g13810),* and *OsGT47-5 (LOC_Os04g32670)* were present in subnetwork 3. This grouping of GT43/GT47 xylan-related genes together with cellulose synthase genes is part of broader co-expression patterns connected with the cell wall. These outputs allow users to evaluate whether selected genes have common expression patterns, significant reciprocal co-expression or possible functional grouping. It is noteworthy to mention that the GAN analysis published by Javaid et al (2024) was performed in RiceFrend platform, which is no longer functioning and lacks many rice genes in its datasets.

**Fig. 6.**
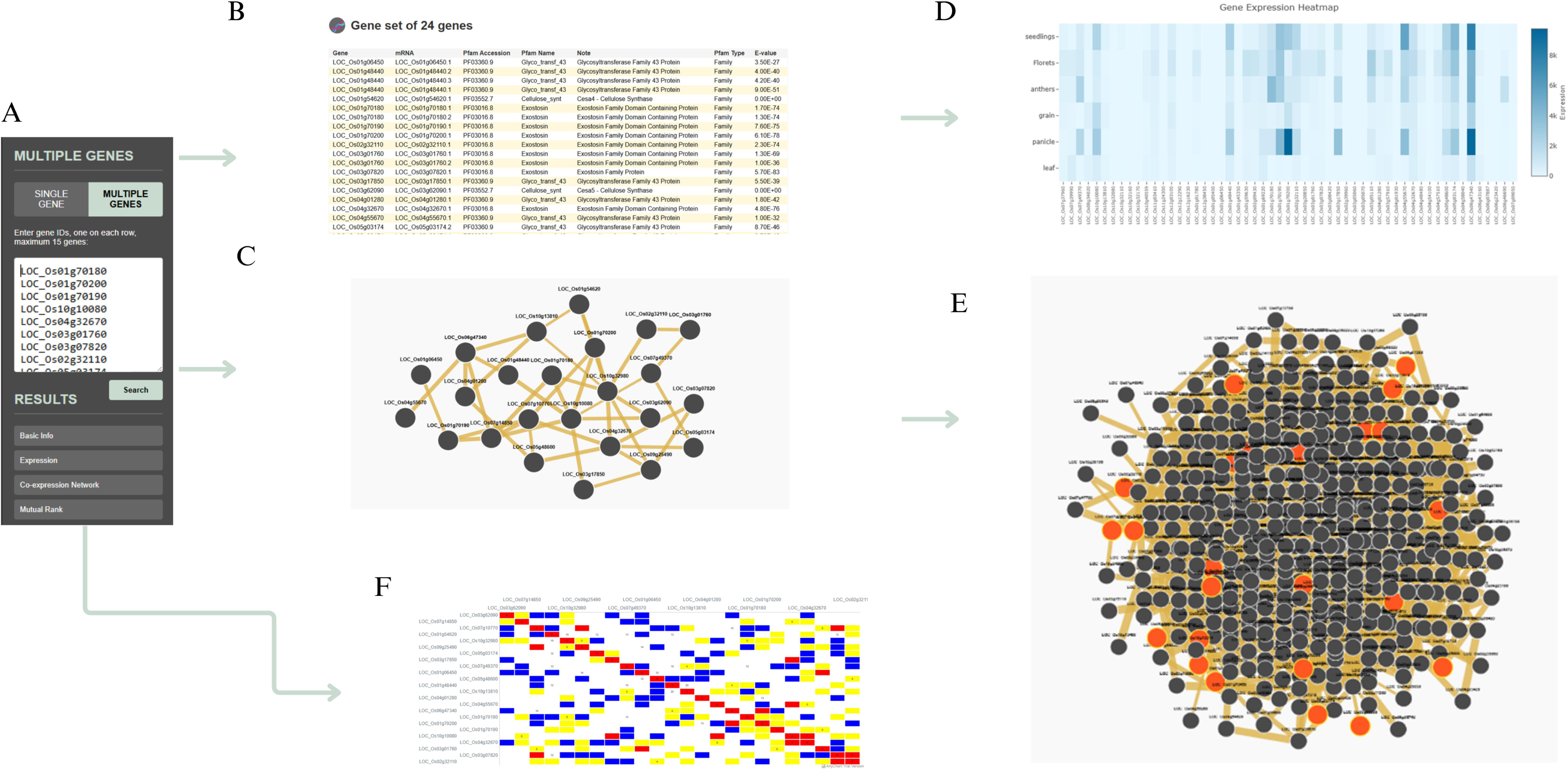
PlantNetX multiple-gene search and co-expression analysis. (A) Interface for multiple gene search in PlantNetX, allowing users to input up to 25 gene IDs. (B) Results panel displaying the gene set with associated gene information, including mRNA ID, Pfam accession, Pfam term name and type, and E-value. (C) Association network generated for the input gene list, illustrating direct connections between strongly coexpressed genes. (D) Gene expression heatmap for the entire input gene set across different tissues and conditions, indicating relative expression levels. (E) A GAN showing the broader connectivity and interactions of the input genes with other coexpressed genes across the rice genome. (F) MR heatmap matrix displaying the pairwise co-expression strength among the queried genes. Lower MR values indicate stronger co-expression. **ALT TEXT:** Six-panel example of a multiple-gene search in PlantNetX. The outputs include the gene-entry interface, annotation table, network among the queried genes, tissue-expression heatmap, expanded GAN, and a MR heatmap showing pairwise co-expression strength.

**Fig. 7.**
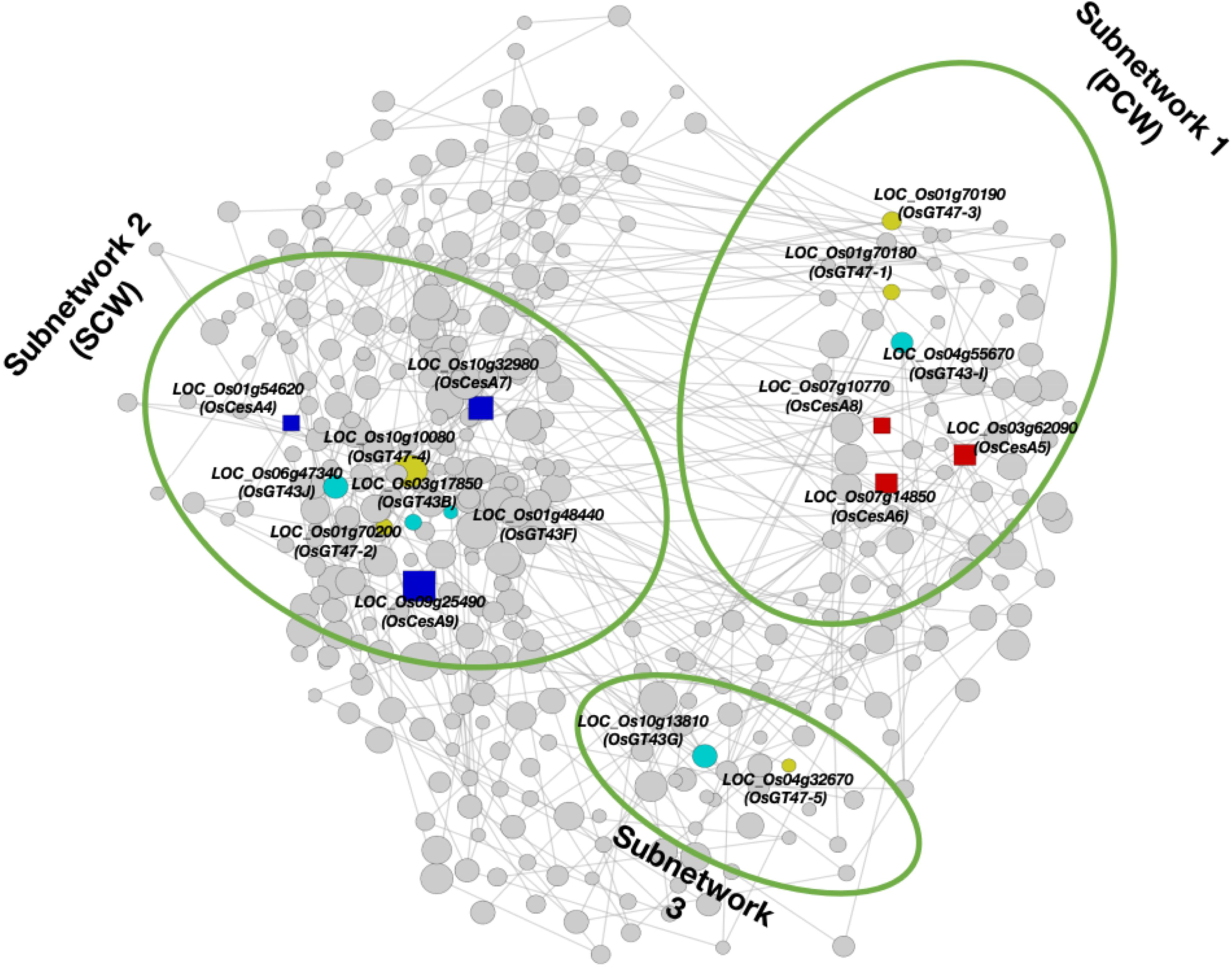
Validation of PlantNetX GAN output using GT43, GT47, and cellulose synthase genes. The RNA-seq-derived GAN separates selected rice GT43 and GT47 glycosyltransferase genes into three subnetworks related to cell wall biosynthesis. Subnetwork 1 includes *OsGT47-3, OsGT47-1,* and *OsGT43I;* Subnetwork 2 includes *OsGT43J, OsGT43B, OsGT43F, OsGT47-4,* and *OsGT47-2;* and Subnetwork 3 includes *OsGT43G and OsGT47-5.* Cellulose synthase genes, including *OsCesA4, OsCesA5, OsCesA6, OsCesA7, OsCesA8, and OsCesA9,* were included as cell wall reference genes to indicate primary and secondary cell wall-related subnetworks. These association patterns are consistent with published evidence for GT43/GT47 interactions and putative xylan synthase complex organization. **ALT TEXT:** GAN selected rice GT43, GT47, and cellulose synthase genes. The network separates the genes into three main subnetworks, with related xylan biosynthesis and cellulose synthase genes grouped according to their co-expression relationships.

### User Interface for Search and Analysis of scRNA-seq datasets

The PlantNetX single cell section enables users to explore gene expression at cell-type level. Users can perform searches using IDs of one or more genes. The outcome includes view summary cards, UMAP-based expression maps, filter outputs by tissue or cell type, compare gene expression overlays, and review cell-type expression tables (Fig. 8). For illustration, *OsCSLD1 (Os10g0578200;* MSU *LOC_Os10g*42750) gene was used because it has been shown experimentally to have a restricted gene expression to rice root hair. *OsCSLD1* encodes a cellulose synthase-like D1 protein needed for rice root hair production and loss-of-function mutants including *rth2* display aberrant root hair formation (Kim *et al*., 2007; Yuo *et al*., 2011). The global UMAP in PlantNetX visualizes the integrated rice cell types across the included scRNA-seq datasets (Fig. 8A). Entering *OsCSLD1* as query in the search box (Fig. 8B) resulted in a summary report with average expression, maximum expression, percentage of expressing cells, detected cell types, and total cells analyzed (Fig. 8C). In the UMAP overlay, expression is primarily observed in cells linked with the root (Fig. 8D), and the cell type expression table indicates the highest expression level was in root hair cells, followed by the epidermis-near-root-hair cells (Fig. 8E). These results are consistent with the published work on *OsCSLD1* (Kim *et al*., 2007; Yuo *et al*., 2011), and support the capacity of PlantNetX to recover expected cell type specific expression patterns. Supplemental Fig. S1 shows detailed tissue-specific cell type expression outputs for *OsCSLD1* in the single gene search.

**Fig. 8.**
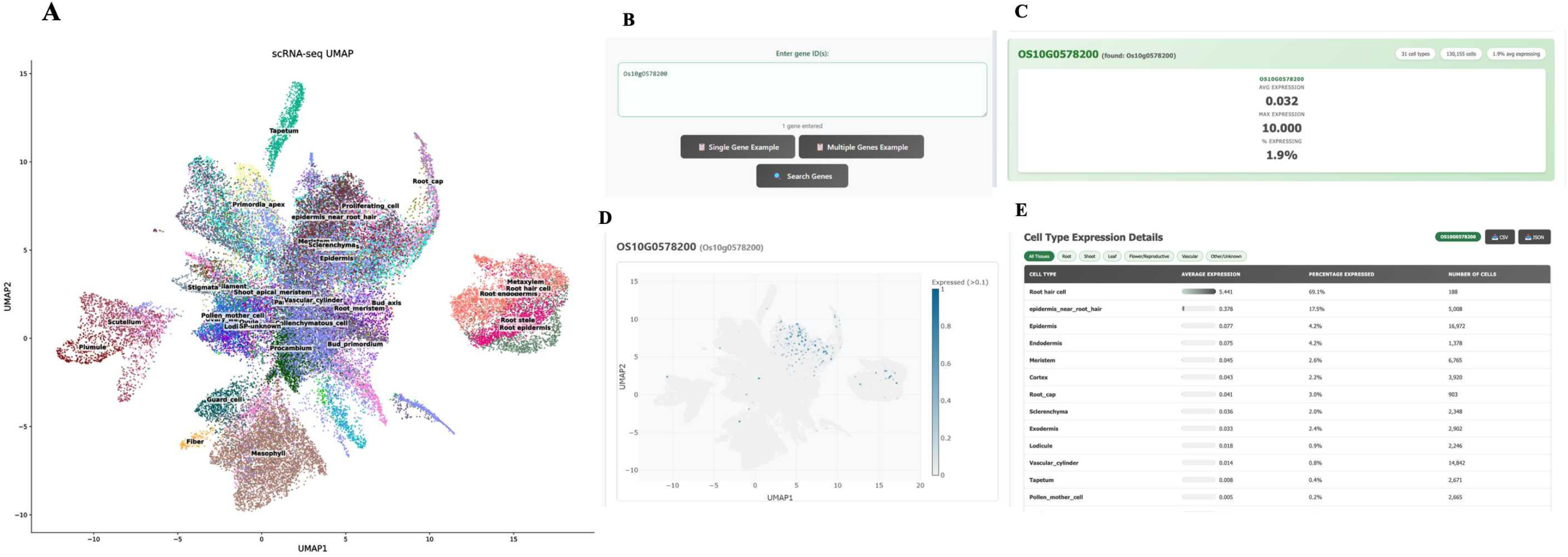
Single-cell validation using *OsCSLD1 (Os10g0578200*) in PlantNetX. (A) Rice single cell clusters annotated on global UMAP. (B) *OsCSLD1* query panel. (C) Summary statistics for *OsCSLD1* comprising average expression, maximum expression, proportion of expressing cells, detected cell types and total cells studied. (D) UMAP overlay of expression of *OsCSLD1* in root-associated cell populations. (E) Cell-type expression table demonstrating enrichment in root hair and epidermis-near-root-hair cell populations. **ALT TEXT:** Five-panel single-cell analysis of OsCSLD1 in PlantNetX. The global UMAP shows annotated rice cell populations, while the expression overlay shows the strongest signal in root-associated cells. The summary table identifies root hair cells as the cell type with the highest expression.

PlantNetX also allows multiple genes to be compared within the same global UMAP coordinate system. For this demonstration, three genes, *Os05g0559600, Os04g0103100* and *Os06g0687900* of GT43 family described by Lee et al. (2014) were selected because they have diverse expression patterns among rice cell types (Lee *et al*., 2014). The global UMAP provides a common coordinate framework for comparing the expression patterns of the queried genes (Fig. 9A), and overlays are shown for individual expression of *Os05g0559600* (Fig. 9B), *Os04g0103100* (Fig. 9C), and *Os06g0687900* (Fig. 9D). The output also contains tables of cell-type expression with average expression, % of expressing cells and # of cells (Fig. 9E). These tables are enabled with tissue-filter buttons so that users can view all tissues or filter to root, shoot, leaf, flower/reproductive, vascular, and other. PlantNetX additionally produces a bar plot of the comparison for all genes searched, allowing direct comparison of the average expression among annotated cell types (Fig. 9F). For instance, *Os05g0559600* is expressed in plumule, root hair cells, endodermis, root stele, epidermis-near-root-hair cells and pollen mother cells with detectable expression. *Os04g0103100* is expressed in endodermis, root cortex, epidermis-near-root-hair cells, exodermis, guard cells, root cap, sclerenchyma and filament. *Os06g0687900* is more highly expressed in root hair cells, epidermis-near-root-hair cells, ovary wall, filament, tapetum, exodermis, endodermis, cortex and shoot apical meristem. These outputs show the efficiency of PlantNetX in comparing gene expression at cellular types. Genes expressed within the same cell types may be prioritized as candidates for shared biological functions or coordinated activity; however, co-expression alone does not establish direct protein–protein interaction or membership in the same protein complex. Supplemental Fig. S2 shows detailed tissue-specific cell type expression outputs for multiple gene example.

**Fig. 9.**
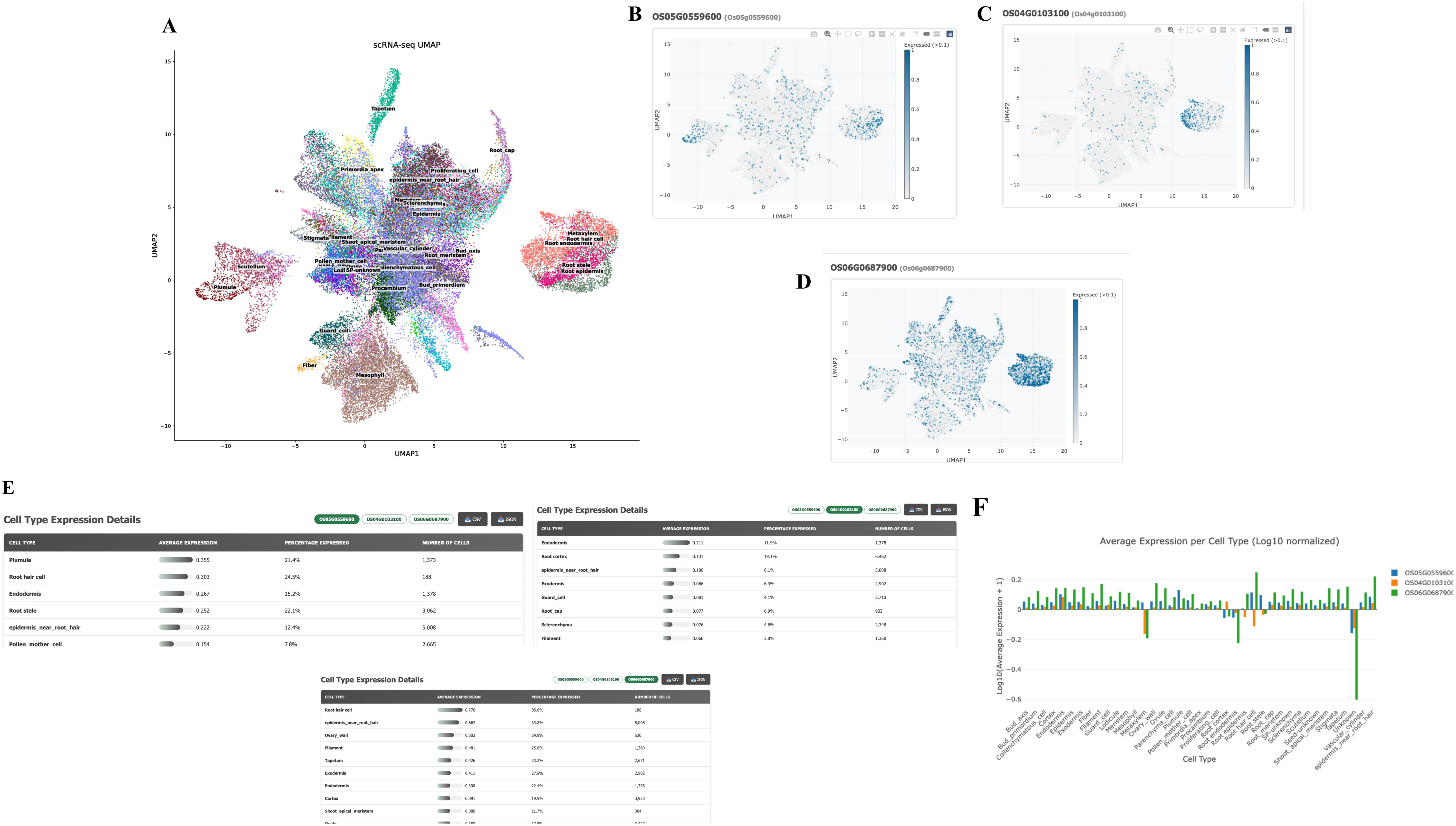
Multiple gene comparison in the PlantNetX single cell module. (A) Global UMAP of integrated and annotated rice single-cell populations used as a common coordinate framework for multi-gene comparison. (B-D) Overlays of UMAP expression for *Os05g0559600, Os04g0103100 and Os06g0687900.* (E) Cell-type expression tables showing average expression, percentage of expressing cells, and assigned number of cells for each cell type, with tissue-filter options for viewing expression across all tissues or selected tissue groups. (F) Bar graph comparing log10 average expression of all queried genes across annotated cell types. **ALT TEXT:** Six-panel comparison of three GT43 genes in the PlantNetX single-cell module. A shared global UMAP is followed by separate expression overlays for each gene, cell-type expression tables, and a bar graph comparing their average expression across annotated cell types.

### Response-time comparison with other plant transcriptomic resources

PlantNetX response time was compared with that of seven representative plant transcriptomic and single-cell databases using the same standardized query conditions. Under the conditions used in this analysis, PlantNetX showed the shortest mean response time at 14.2 s. The mean response times of the other resources ranged from 18.7 s for PPRD to 35.8 s for PlantscRNAdb **(Fig. 10)**. These results suggest that the precomputed data structure and query-based retrieval system used in PlantNetX support rapid access to gene expression and network outputs.

**Fig. 10.**
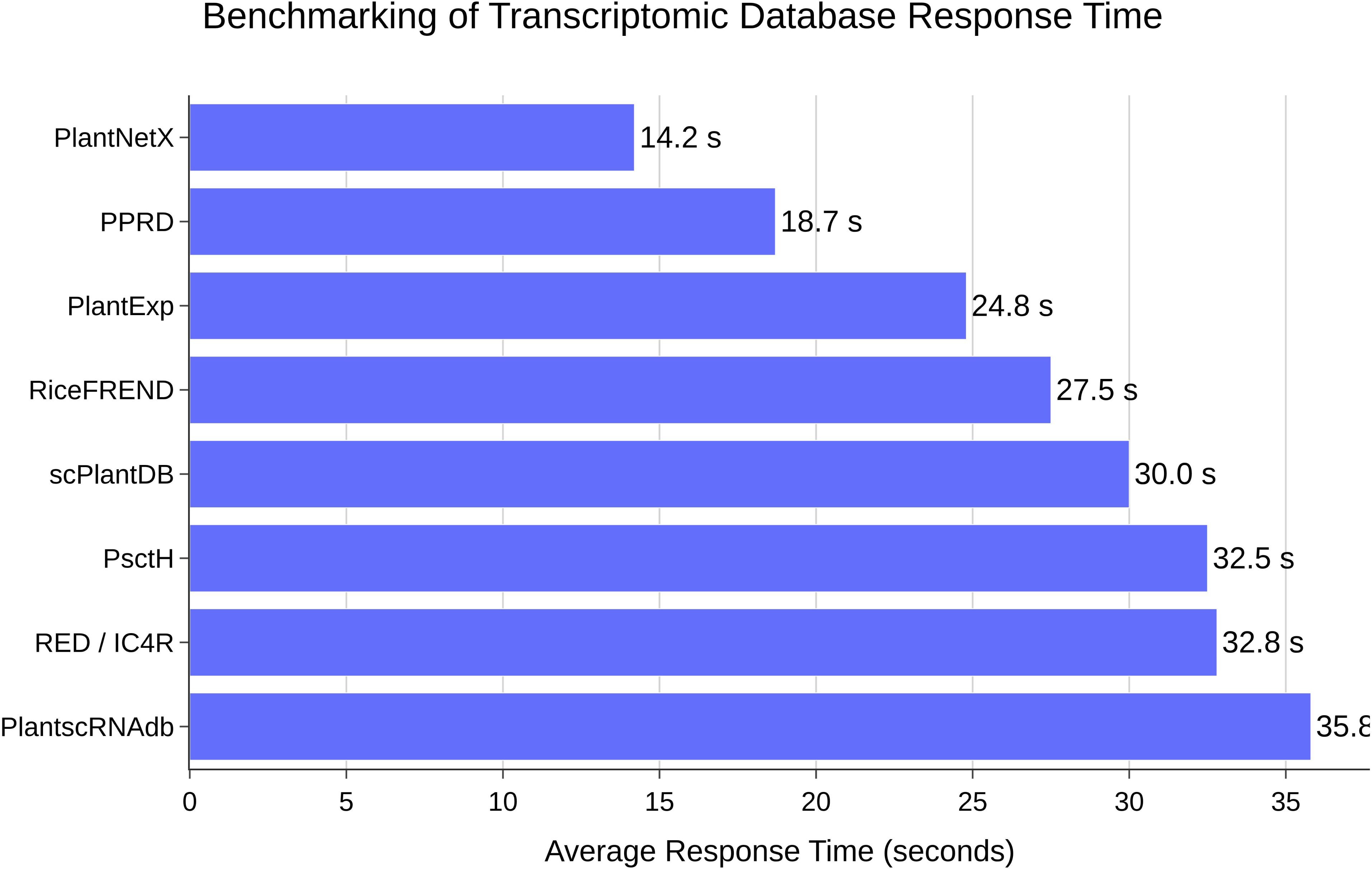
Response-time comparison of PlantNetX with representative plant transcriptomic and single-cell databases. Response time was measured from submission of a standardized gene query until the principal results page was fully displayed. Each platform was tested in 20 independent runs using the same computer, browser, internet connection, query gene, and testing period. Bars represent the mean response time across the 20 runs. Lower values indicate faster response under the conditions used in this benchmark. **ALT TEXT:** Bar graph comparing the mean response times of PlantNetX and seven plant transcriptomic or single-cell databases. PlantNetX has the shortest mean response time at 14.2 seconds, whereas PlantscRNAdb has the longest at 35.8 seconds. Each value is based on 20 independent technical runs.

## Discussion

PlantNetX is developed as an expandable platform for plant transcriptome analysis. Rice was selected as the first model system because of its well annotated genome, availability of large public transcriptomic datasets, and importance as a crop and monocot model. Yet, PlantNetX platform intends to go beyond rice by include other plant species, tissue-specific RNA-seq datasets, single-cell datasets, and better cell-type-level expression sections. PlantNetX allows linking expression profiles to tissue-specific GANs, candidate gene ranking and single-cell interpretation because both RNA-seq and scRNA-seq datasets are present in the same platform. While other platforms such as PlantExp, (Liu *et al*., 2023) and PPRD (Yu *et al*., 2022) are useful for expression browsing, they do not provide integrated GAN analysis or scRNA-seq-based cell-type interpretation. PlantNetX also incorporates recently available rice RNA-seq and scRNA-seq datasets that are not consistently represented in existing resources, providing broader and more current transcriptomic coverage.

Network-based techniques like GAN are growing into key tools for crop development, as these approaches assist relate transcriptome variance to biological function. Rather than evaluating individual genes in separate groups, GAN or weighted gene co-expression network analysis (WGCNA) based approaches cluster genes into co-expression modules that are potentially common pathways, regulatory programs or trait-associated activities. Genes with similar expression patterns tend to be involved in similar pathways, same regulatory programs or coordinated cellular processes (Serin *et al*., 2016; Usadel *et al*., 2009). These techniques were utilized to uncover nitrogen-responsive gene modules and putative regulators related to nitrogen-use efficiency in rice, such as transcription factors, transporters, kinases, and other regulatory genes (Sharma *et al*., 2023). Co-expression research has also shown key modules and hub genes linked to cold-stress response and recovery, including as genes involved in autophagy, sugar metabolism, and stress adaptation (Zeng *et al*., 2022). In drought and heat stress response research, similar network methodologies have been employed, and WGCNA has been used to uncover core stress-responsive genes and biological processes associated with stress tolerance (Cao *et al*., 2024). These examples demonstrate the ability of network-based analysis to reduce massive transcriptome datasets to selected candidate genes and modules for downstream validation, breeding and crop improvement. In this sense, PlantNetX platform offers a practical framework for plant biotechnology to find candidate genes, explore tissue-specific associations, compare gene families, and select targets for downstream validation (i.e., mutant analysis, overexpression, CRISPR/*Cas*9 editing or protein-protein interaction testing).

One of the main strengths of PlantNetX is that it goes beyond a simple gene expression lookup. The platform combines gene annotation, RNA-seq expression, GAN analysis, MR-based co-expression ranking and single-cell expression visualization into a unified process. It also supports both single-gene and multiple-gene searches, allowing users to examine individual candidates or compare gene families and pathway-related gene sets within the same workflow. The RNA-seq based design has an edge over the older rice co-expression resources. For example, although RiceFREND proved to be helpful for the retrieval of co-expressed genes in rice, it was established using large-scale microarray expression data (Sato *et al*., 2013; Zainal-Abidin *et al*., 2022), which lacks resolution and sensitivity and have 24,448 MSU rice gene loci (some rice genes are not represented in the datasets). Furthermore, RiceFREND platform is no longer functioning (as of May 2026). In contrast, PlantNetX uses RNA-seq datasets, which can be processed uniformly with higher resolution to find expression variance across tissues and biological circumstances, and almost all 55,801 MSU gene loci are represented in RNA-seq datasets. In addition, RiceFREND platform has practical limitations on the visualization of large networks. For example, if the number of nodes exceeds 100, only the downloadable file is provided, and if the number of nodes exceeds 300, neither the network display nor the Cytoscape (Franz *et al*., 2016) upload file is available. PlantNetX provides MR-threshold-based network exploration, interactive Cytoscape.js visualization, and downloadable outputs; however, the practical size of an interactive network depends on browser performance, the number of nodes and edges, and the user’s computing resources. Furthermore, PlantNetX uses MR to improve linkages based on Pearson’s correlation, a commonly used method among plant co-expression resources to identify biologically significant gene relationships. The MR is an essential metric in GAN analysis since it measures the reciprocal strength of connection between two genes. MR-based ranking has been commonly used in plant co-expression resources for improving the prioritization of candidate genes (Obayashi *et al*., 2022; Sato *et al*., 2013). A lower MR value suggests a higher reciprocal co-expression connection between the two genes. A stringent MR cut-off results in a smaller, more particular network. A greater MR cut-off results in a larger network with more nodes and edges, but possibly weaker association. PlantNetX thus enables the user to adjust the MR threshold based on his analytical goal, either to facilitate the focused selection of potential genes or to explore the network in a more general sense. PlantNetX further enhances usability using Cytoscape.JavaScript based visualization with interactive node and edge exploration, zooming, layout tweaking and export compatible results. This enables users to go more swiftly from gene searching to network understanding.

The single-cell component provides PlantNetX platform with cellular resolution information. Several plant scRNA-seq resources have been developed, including PlantscRNAdb (Chen *et al*., 2021) for plant marker genes and cell-type information, PsctH for exploring plant single-cell transcriptome landscapes, and scPlantDB (He *et al*., 2024) for marker exploration, cell-type comparison, visualization and data download. However, these resources generally focus on marker searching, cell-type annotation or single-cell visualization, and are not directly related to GAN analysis based on RNA-seq data. Some may not contain full coverage of rice tissue or do not include the most recently publicly available rice scRNA-seq datasets. PlantNetX now hosts 9 up to date rice scRNA-seq datasets, including 81 samples. PlantNetX integrates these data with RNA-seq generated GAN results to give users a single platform to assess candidate genes at tissue- and cell-type-resolution. Future, versions of PlantNetX will contain tissue-specific sections, such as root, leaf, reproductive tissue, seedling, floret, spikelet and meristem. For each segment, cell-type composition, UMAP views, gene search results, and cell-type expression summaries will be provided separately. This should help links tissue-specific GANs to matched cell-type-level expression for more accurate assessment of potential genes. The platform is freely accessible without registration, and users can download gene-expression profiles, co-expression results, network outputs, and single-cell expression summaries for downstream analysis.

Under the standardized testing conditions used in this study, PlantNetX showed a shorter mean response time than the other evaluated plant transcriptomic and single-cell resources (Fig. 10). This result suggests that the use of precomputed expression matrices, co-expression outputs, and query-specific data retrieval supports efficient access to the platform’s main analytical features. Because response time can vary with server load, internet connection, geographic location, browser configuration, and query complexity, these findings should be interpreted within the conditions of the present comparison. Overall, the benchmark suggests that PlantNetX provides a responsive and practical interface for transcriptomic data exploration.

PlantNetX has, however, certain restrictions. The accuracy of GAN is dependent on the quality, diversity and tissue coverage of the available RNA-seq datasets and some developmental stages, stress situations or genotypes may be underrepresented. Co-expression can indicate a potential functional association, but it does not demonstrate direct regulation or physical interaction between proteins. These relationships require independent experimental validation. Similarly, the single-cell section depends on the quality and consistency of public scRNA-seq datasets, and rare, transitional or tissue-specific cell types may require additional refining as new datasets become available. PlantNetX is intended to be a dynamic and extensible resource. In the future, the platform will incorporate new RNA-seq and scRNA-seq datasets routinely as they become available. Other capabilities will include stress-specific analysis, gene ID conversion, user-uploaded gene lists, ortholog search, cross-species comparison, and integration with additional omics layers. Future updates will include built-in conversion between MSU LOC and RAP-DB identifiers. With the addition of more datasets and plant species, PlantNetX can transition from a rice-centred platform to a larger plant transcriptome resource for functional genomics and crop development.

PlantNetX enables plant functional genomics by connecting transcriptome relationships to biological context. The tool lets researchers to prioritize enormous lists of genes and explore their activity in tissues and cell types. This can lead downstream research such as mutant screening, CRISPR/Cas9 editing, overexpression analysis and protein-interaction testing. PlantNetX can be valuable for future crop development studies.

## Conclusion

PlantNetX integrates bulk RNA-seq and single-cell RNA-seq data into a single platform to facilitate the study of gene relationships at both the tissue and cell-type levels. Users can find co-expressed genes, explore MR-based networks, compare expression patterns and download data for additional study. These features can be useful for reducing big gene lists to more compact candidates for experimental testing. The platform may therefore be valuable in investigations of plant growth, stress responses, nutrient usage, yield, cell-wall biosynthesis and other agronomic aspects. PlantNetX currently focuses on rice, but its architecture will allow the addition of new datasets, plant species and analytical tools in the future. Future improvements will include enhanced tissue-specific networks, updated cell-type annotations, gene ID conversion, user supplied gene lists, ortholog searches and cross-species comparisons. PlantNetX provides a viable approach to connect big transcriptome datasets to biological concerns and to facilitate gene identification and research in crops.

## Abbreviations

AJAX: asynchronous JavaScript and XML
CSS: Cascading Style Sheets
CSV: comma-separated values
GAN: Gene Association Network
GEO: Gene Expression Omnibus
GO: Gene Ontology
GTF: Gene Transfer Format
HTML: HyperText Markup Language
JSON: JavaScript Object Notation
MR: Mutual Rank
PCA: principal component analysis
PCC: Pearson correlation coefficient
RNA-seq: RNA sequencing
RPK: reads per kilobase
scRNA-seq: single-cell RNA sequencing
SRA: Sequence Read Archive
TPM: transcripts per million
UMAP: Uniform Manifold Approximation and Projection
UMI: unique molecular identifier.

## Supplementary data

**Table S1.** Curated bulk RNA-seq datasets incorporated into PlantNetX, including accession, sample, tissue, experimental description, and data-source information.

**Table S2.** Rice single-cell RNA-seq datasets incorporated into PlantNetX, including accession and tissue-source information.

**Fig. S1.** Tissue-filtered cell-type expression profiles of *OsCSLD1* across the PlantNetX single-cell datasets.

**Fig. S2.** Tissue-filtered cell-type expression profiles for the three-gene comparison used in the PlantNetX single-cell analysis.

## Acknowledgment

We thank the Ohio Supercomputer Center for computing resources and Ohio University OIT for hosting PlantNetX. This work was supported in part by the Ohio University Baker Fund Award and Student Enhancement Award.

## Author Contributions

M.A.N., S.N., and A.F. conceived and designed the study. M.N. and S.N. contributed equally to dataset collection, data organization, data analysis, PlantNetX platform development, and manuscript preparation. A.F. contributed to study design, supervision, interpretation of results, manuscript revision, and overall project guidance. All authors reviewed and approved the final manuscript.

## Data Availability

The publicly available bulk RNA-seq and scRNA-seq datasets analysed in this study are deposited in the NCBI GEO and SRA under the accession numbers listed in Supplementary Tables S1 and S2. The curated metadata, processed expression matrices, Pearson correlation and MR outputs, single-cell expression summaries, marker-gene information, source code, and analysis scripts generated for PlantNetX are openly available in the PlantNetX GitHub repository at https://github.com/Mohsin-OU/PlantNetx. The PlantNetX web resource is freely accessible without registration at https://plantnetx.academic.kube.ohio.edu/. No novel biological materials were generated in this study.

## Supplementary Material

**Supplementary Table S1. Bulk RNA-seq datasets integrated into PlantNetX.** This table provides the curated bulk RNA-seq datasets used for PlantNetX RNA-seq expression analysis and GAN construction. It includes dataset accession information, sample details, tissue or organ type, experimental description, and dataset source.

**Supplementary Table S2. Single-cell RNA-seq datasets integrated into PlantNetX.** This table provides the curated rice scRNA-seq datasets used for PlantNetX single-cell analysis. It includes dataset accession information, and tissue source.

**Supplementary Fig. S1.** Tissue-wise cell-type expression distribution of *OsCSLD1* in PlantNetX. Cell-type expression tables for *OsCSLD1* showing average expression, percentage of expressing cells, and number of cells across tissue-filtered categories, including root, shoot, leaf, flower/reproductive, vascular, and other/unknown groups. The tissue-wise view shows that *OsCSLD1* expression is enriched mainly in root hair and epidermis-near-root-hair cell populations, consistent with its role in rice root hair development.

**ALT TEXT:** Multiple cell-type expression tables showing OsCSLD1 expression across root, shoot, leaf, reproductive, vascular, and other tissue groups. The highest expression is observed in root hair and epidermis-near-root-hair cell populations, while expression is lower in most other cell types.

**Supplementary Fig. S2.** Tissue-filtered multiple-gene expression output in PlantNetX. (A) Multiple-gene search interface showing the queried genes *Os05g0559600, Os04g0103100,* and *Os06g0687900*, with summary displaying average expression, maximum expression, percentage of expressing cells, total detected cell types, and total cells analyzed. (B - D) Tissue-filtered cell-type expression tables showing average expression, percentage of expressing cells, and number of cells for the queried genes across root, shoot, leaf, flower/reproductive, vascular, and other/unknown tissue groups.

**ALT TEXT:** Four-panel figure showing a multiple-gene query for *Os05g0559600, Os04g0103100,* and *Os06g0687900* and their cell-type-specific expression across tissue groups. The first panel presents the query and summary statistics, while the remaining panels compare average expression, percentage of expressing cells, and cell numbers across root, shoot, leaf, reproductive, vascular, and other cell populations.

